# Mitochondrial genetics of exceptional longevity in multigeneration matrilineages

**DOI:** 10.1101/361899

**Authors:** Richard A. Kerber, Elizabeth O’Brien, Ron Munger, Ken R. Smith, Richard M. Cawthon

## Abstract

Some heritable mitochondrial DNA (mtDNA) sequence variants may slow the rate of aging. The European mitochondrial haplogroup K has previously been reported to be increased in frequency in centenarians and nonagenarians relative to its frequency in younger individuals, by standard case/control study designs. To select for mitochondrial genomes likely to carry beneficial genetic variants, we screened a large genealogical database (the Utah Population Database, UPDB) for mitochondrial lineages in which the frequency of survival past 90 years was significantly higher than in the general population, and also significantly higher than in close non-matrilineal relatives. We ranked 14,900 distinct matrilineages by the strength of their association with longevity. Full sequencing of the mtDNAs from a single individual from each of 53 matrilineages in the top longevity ranks and each of 374 control matrilineages from the general Utah population, followed by analyses of the mtDNA haplogroup frequencies, identified haplogroup K2 as the haplogroup most enriched in frequency by the longevity selection (Odds Ratio = 23.05). We then analyzed overall survival and cause-specific mortality in the several thousand individuals aged 40 years or older whose mtDNA genotypes could be imputed from the 374 fully sequenced control mtDNAs. In these control matrilineages Haplogroup K2 individuals (n=332) enjoyed a significantly lower all-cause mortality risk than the general population (HR=0.81), attributable in part to a significantly lower risk of dying from heart disease (HR=0.50), as well as lower (though not significantly lower) risks of dying from cancer (HR=0.72) and diabetes (HR=0.74). Furthermore, K2 was the only haplogroup in which mortality was reduced for all three of these common causes of death.

## Introduction

The heritable component to a long and healthy life is likely to involve the actions and interactions of both nuclear and mitochondrial genetic variants. Mitochondrial haplogroup K has been reported to be increased in frequency in exceptionally long-lived individuals as compared to younger controls, in French [1], Irish [2], and Finnish [3] populations. Other studies, also with a case-control design, have reported an association of haplogroup K with decreased risk of Parkinson Disease [4], Alzheimer Disease in APOE 4 allele carriers [5], pancreatic cancer [6], thyroid cancer [7], and both transient ischemic attack and ischemic stroke [8].

Since the mtDNA is always maternally transmitted, all members of a multigeneration matrilineage share the same mtDNA sequence; therefore, one need only sequence the mtDNA of one individual in a matrilineage in order to know the mtDNA sequence of all members of the matrilineage. Using the Utah Population Database (https://healthcare.utah.edu/huntsmancancerinstitute/research/updb/), a large genealogical database with records on more than 8 million individuals, we identified the matrilineages with the highest frequency of exceptionally long-lived individuals, and collected blood samples from 53 of them for full mtDNA sequencing. These longevity-selected mtDNAs were compared to mtDNAs from 101 different matrilineages from the Utah CEPH (Centre d’Etudes du Polymorphisme Humain) families [9], and from an additional 273 distinct matrilineages from the Cache County Memory Study [10] provided by Dr. Munger, to identify mitochondrial haplogroups and mtDNA sequence variants enriched in the longevity-selected matrilineages. Haplogroups and sequence variants found in the control mtDNAs were also tested for association with all-cause, heart disease, cancer, and diabetes mortality among all of the known control matrilineage members who survived to 40 years or older, several thousand individuals.

## Results

The two tables below present mtDNA haplogroup and mtDNA sequence variant associations with membership in a set of 53 Utah matrilineages selected for the frequent occurrence of extreme longevity (LSL, or Longevity Selected Lineages); overall survival beyond age 40 years in 374 control matrilineages from the general Utah population (273 from the NIA-funded Cache County Study on Memory Health and Aging, and 101 from the Utah CEPH families); and mortality from heart disease, cancer, and diabetes in those 374 matrilineages. Haplogroup K subclade K2, and the T9716C sequence variant shared by all haplogroup K2 members, showed significantly lower all-cause and heart disease mortality, as well as reduced (though non-significantly) risks of dying from cancer and diabetes.

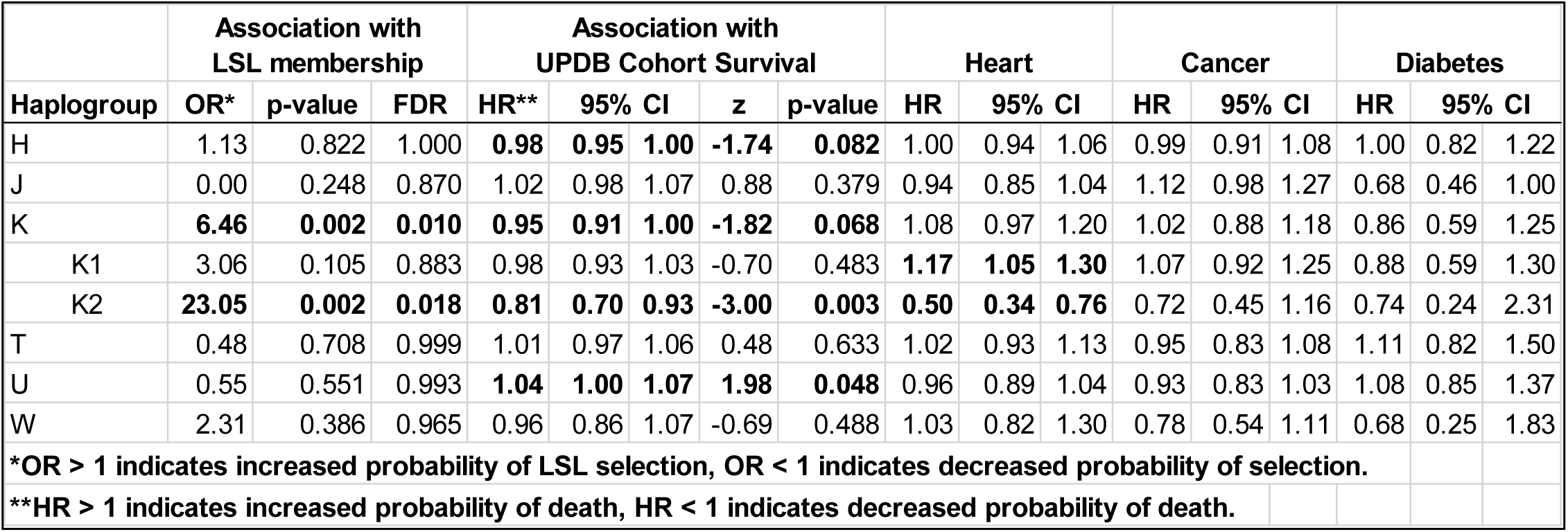

Overrepresentation among the 53 LSL (longevity-selected) matrilineages and association with reduced mortality in the UPDB cohort (controls) are independent tests, and represent a replication of results (albeit within the same population) for the mtDNA nucleotide changes at 9716, 16224, and 146, and is highly suggestive for 10550, 3480, 9055, 14167, 11299, 9698, and 523.1/523.2. Some variants found in the longevity-selected matrilineages did not occur in the control matrilineages, and so could not be tested against survival in the control lineages.

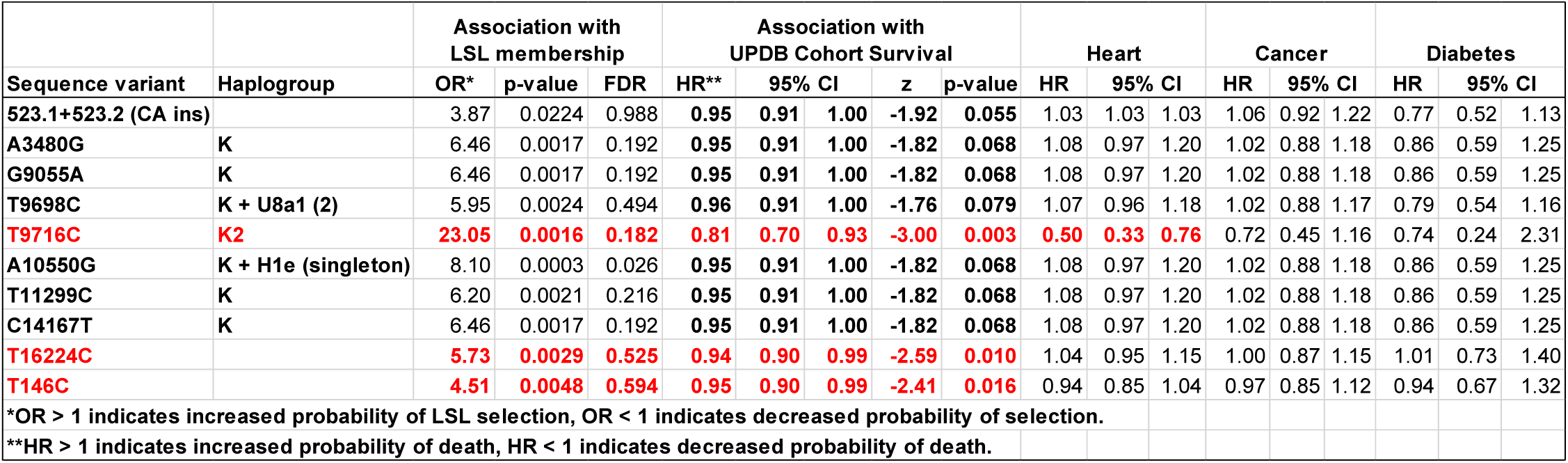

## Discussion

To our knowledge this report is the first to use survival and cause-specific mortality data in multigeneration matrilineages to identify mitochondrial genetic variants that may slow aging and contribute to longevity by lowering the risk of not just one, but several common causes of death. The previously reported published case-control studies [1–8] attributing protection from several aging-related diseases to haplogroup K, combined with the evidence presented here of haplogroup K2’s broad health benefits, based on morbidity and mortality data from multiple multigeneration matrilineages, strongly support the conclusion that haplogroup K, and especially the K2 subclade, contributes to better survival by reducing the risk of dying from not just one, but multiple common diseases. How haplogroup K2 accomplishes this has yet to be determined, and may involve multiple mechanisms.

Reactive Oxygen Species (ROS) are generated when electrons fall off the Electron Transport Chain (ETC) prematurely (i.e. at any point prior to their donation to oxygen by Complex IV to make water). The steeper the hydrogen ion gradient across the inner mitochondrial membrane, the harder it is for electron transport to be coupled to the pumping of protons up their gradient, and the more likely it becomes for electrons to fall off the ETC prematurely. The free radical theory of aging argues that ROS damage to DNA, proteins, and lipids accumulates with age, eventually causing cell and tissue dysfunction, disease, and death.

Mitochondrially encoded amino acid substitutions in the proteins of oxidative phosphorylation could lower the rate of ROS production per unit oxygen metabolized in at least three different ways: 1) making the hydrogen ion gradient less steep by increasing the leak of protons, without ATP production, across Complex V; 2) making the hydrogen ion gradient less steep by decreasing the number of protons translocated per electron transported down the full ETC; or 3) for a given level of hydrogen ion gradient, increasing the efficiency of electon transport through the ETC, so that fewer electrons fall off prematurely.

Because haplogroup K was absent in endurance athletes (present in 0 of 52 endurance athletes vs. 48 of 1060 controls) and associated with longevity, Niemi et al. [11] suggested that it may be an “uncoupling genome.” Uncoupling could be due to the G9055A-encoded amino acid substitution (Ala->Thr) in ATP synthase subunit 6 promoting proton leakage through Complex V [12]. This would be expected to reduce the efficiency of oxidative phosphorylation (limiting athletic performance) and decrease the production of reactive oxygen species (extending lifespan). Moreover, an uncoupling mitochondrial genome may prevent or reduce the rise in ROS levels associated with excessive caloric intake, thereby reducing the risk of developing insulin resistance and diabetes. Whether cells from haplogroup K2 individuals actually have a decreased mitochondrial membrane potential and lowered rates of mitochondrial ROS production remains to be tested.

While the G9055A variant is shared by all haplogroup K carriers, the K2 subclade appears to have health benefits above and beyond those shared by K haplogroup members generally. Therefore, one or more sequence variants found in K2 but missing from other K subclades may further promote health. The T9716C sequence variant is specific to K2, shared by all K2 individuals, and not found in any other mitochondrial haplogroups. Interestingly, it is a synonymous nucleotide substitution, i.e. it does not alter the amino acid sequence of its encoded protein.

It has been reported that species with higher maximum lifespans tend to have higher GC-content in their mtDNA sequences [13]. Higher GC-content may protect mitochondrial genomes, since single-stranded DNA is more prone to damage of various types than double-stranded DNA, and stretches of DNA with higher GC-content will spend less time open and single-stranded than otherwise similar sequences with lower GC-content, especially at high temperatures. Slippage of DNA polymerases during DNA replication, resulting in small deletions and insertions, also occurs more frequently at higher temperatures, and more frequently in AT-rich sequences than in GC-rich sequences. A recent study showed that in human cells mitochondrial physiology maintains mitochondria at a temperature near 50°C [14], much higher than the typical core body temperature of around 37°C, making the hypothesis of differences in mtDNA damage rates being sensitive to haplogroup differences in GC-content even more attractive. Among European mitochondrial haplogroups, haplogroup K is among the highest for GC content. In haplogroup K2 the bolded red “t” at position 9716 in the revised Cambridge Reference Sequence, 5’-attttactggg**t**ctctattttac-3’, is mutated to a “c”. Perhaps the AT-rich 7-base direct repeat (attttac) flanking 9716 makes this region prone to replication errors, and the increased GC-content provided by T9716C provides increased stability, lowering error rates. The just upstream T9698C base change, which is specific to, and shared by all, K haplogroup members, further raises GC-content, perhaps further stabilizing the region during replication. Further work will be needed to investigate these and other possible explanations for haplogroup K2’s health benefits.

## Research subjects & methods

### Longevity-selected mitochondrial lineages

The Utah Population Database (UPDB) is a genealogical database with more than 8 million family history, demographic, and medical records of Utah residents. This is primarily a Caucasian population of Northern European origin.

We identified 14,900 human mitochondrial lineages in the UPDB with female founders born from 1800–1860, and selected those matrilineages in which the proportion of members over 65 who survived to age 90 was significantly greater than the proportion of the general population over 65 who survived to 90. We further selected for matrilineages whose members showed significantly better survival than their non-matrilineal relatives. 53 matrilineages resulting from this filtering process were selected for further study. One blood sample from an individual age 65 years or older was obtained from each of the 53 matrilineages.

### Control mitochondrial lineages

One hundred one individuals from different matrilineages (based on UPDB identification of the female founders) were chosen from the 47 three-generation Utah CEPH (Centre d’etude du polymorphisme humain) families [12] for full mtDNA sequencing. The full mtDNA sequences of 273 individuals from the Cache County Memory Study [13], who were determined to be from an additional distinct set of UPDB matrilineages, were provided by Dr. Munger.

### DNA sequencing

The longevity-selected mtDNAs and the Cache County mtDNAs were fully sequenced by Family Tree DNA (https://www.familytreedna.com/), Houston, TX. Overlapping PCR products covering the entire mtDNA sequence were prepared, using primer sets that have been proven not to amplify from mitochondrial-like pseudogenes in the nuclear DNA (a.k.a. numts), and sequenced by standard Sanger sequencing. Some of the Utah CEPH mtDNAs were sequenced by Family Tree DNA, and others were sequenced by the DNA sequencing core laboratory at the University of Utah also using mtDNA-specific primers.

### Genetic association, survival, and cause-specific mortality analyses

Survival was analyzed in Utah Population Database cohorts who were alive and followed from the time they turned 40 years old, until death, loss to follow-up, or present time. Overall survival and deaths from specific causes were compared to mitochondrial genetic variation measured as haplogroup, subclade, and specific variant. Full mitochondrial genotypes were extrapolated to all members of each genotyped person’s matrilineage. Analyses were adjusted for sex, year of birth, connectedness to pedigree structures, nuclear familial excess longevity, and familial standardized mortality ratios for specific causes of death.

## Acknowledgements

We thank Dr. Mark Leppert for providing DNA samples from 101 Utah CEPH individuals for full mtDNA sequencing, Missy Dixon for Utah CEPH grandparent survival data, and Alison Fraser and the UPDB staff for survival and cause-specific mortality data for all Utah CEPH and Cache County matrilineage members. We thank the Pedigree and Population Resource (funded by the Huntsman Cancer Foundation) for its valuable role in the ongoing collection, maintenance, and support of the Utah Population Database.

